# Different, same, or arrhythmic: an empirically informed benchmark of differential rhythmicity methods

**DOI:** 10.64898/2026.10.21.760649

**Authors:** Bharath Ananthasubramaniam

## Abstract

Circadian rhythms represent a key biological dynamic across all kingdoms of life. These cell-autonomous rhythms generated by a gene-regulatory network regulate processes in a time-of-day-specific manner primarily via transcriptional rhythms in a significant fraction of genes. The controlled experiment paradigm can be used to study causal effects at the molecular level using transcriptomic RNA-seq data even in this dynamic scenario. Multiple tools for the analysis of such data, termed differential rhythmicity (DR) analysis, are available but guidance on method choice remains limited. Here, we present a systematic benchmarking study of 12 different methods from 5 packages using a synthetic model of RNA-seq count data that incorporates both empirically estimated mean-dispersion trends and concomitant differential expression changes. By linking method performance to experimental conditions and analytical goals, we provide practical and simple rules of thumb that will help biologists choose appropriate DR methods and extract reliable biological insight from time series RNA-seq data.

## Introduction

Approximately 24-hour “circadian” variation in physiology to adapt to day-night environmental changes on Earth is fundamental to animals, plants, and fungi. These cell-autonomous molecular circadian rhythms are generated by a gene-regulatory network, *the core clock*, in eukaryotes. The time-of-day information encoded by the clock is distributed to most other physiological processes by transcription factors (TFs) that are part of the core clock. The TFs regulate the expression of clock-controlled genes genome-wide that are the proximal output of *the clock*. Therefore, transcriptomics is an essential tool in the study of circadian clocks, and high-throughput controlled experiments evaluate concordance and discordance between rhythmic expression across conditions to gain mechanistic insight into clock inputs and outputs.

Tools for such differential rhythmicity (DR) analysis, the sister of differential expression (DE) analysis for static phenomena, have proliferated in recent years, which has increasingly left users wondering “which method should I use for my study?” Some studies have investigated the shortcomings of DR analysis techniques too. We identified a statistical shortcoming in the intuitive precursor to DR analysis based on Venn diagrams that results in an overestimation of the number of DR genes [1]. At the same time, there is both little overlap in rhythmic genes in the same wild-type tissue under similar conditions across different labs and studies [2] and a lack of consistency in the output of different tools applied to the same dataset [3]. However, it remains unclear whether this is a reflection of the limitations of the tools, the underlying biological reality, or some combination of the two.

Here, we present a systematic benchmarking study providing practical guidance linking DR method choice to RNA-sequencing experimental conditions. We consider in particular two previously overlooked complexities of the DR analysis problem: the correlation between mean and dispersion of the count data, a key feature of RNA-seq data, and the confounding of DR analysis by concurrent DE, which is often present in controlled experiments. We evaluated methods under realistic, user-relevant conditions using synthetic data with ground truth and stratified our insights to provide structured, evidence-based rules of thumb for method selection. By turning methodological benchmarking into practical rules of thumb, this work aims to improve the reliability with which time series RNA-seq data are translated into biological insight.

## Methods

### Data-Generating Model

To compare different methods, we needed datasets where the ground truth was known and controllable. We therefore expanded the approach from Soneson and Delorenzi [4] to generate time series RNA-seq count data with the following properties:

- The data are drawn from a negative binomial (NB) distribution with an empirically estimated mean-dispersion trend, which is an essential characteristic of RNA-seq data.
- The time series are generated under two different conditions at the same time points with differential expression in a random subset of genes.
- The data included a tunable fraction of rhythmic genes with known amplitudes and phases in the two conditions. This resulted in genes with “gain”, “loss,” “same” and altered (“change”) rhythms between the conditions.

The *G* genes in the dataset were defined at *t*_*n*_, *n* = 1, …, *N* under each of the two conditions, control (*ctrl*) and intervention (*expt*). A fraction *f*_DE_ and number *G*_DE_ of the genes were differentially expressed (DE). Further, a fraction *f*_R_ and number *G*_R_ of the genes were rhythmic in at least one condition. The extent of differential expression was modeled as *η*_*g*_ *~ η*_min_ + Exp(*η*_spread_) and *η*_*g*_ = 0 for genes without differential expression.

The numbers of genes across the four rhythmic groups (same, gain, loss, change) were drawn from a multinomial distribution with a uniform prior such that *G*_R_ = *G*_same_ + *G*_gain_ + *G*_loss_ + *G*_change_. The rhythm parameters for genes are drawn such that *A*_*g*_ *~ a*_min_ + Exp(*a*_spread_), *ϕ ~ U* [0, 2*π*), where *A*_*g*_ is log_2_ fold-amplitude relative to mesor. The parameters in the four groups are further constrained as follows: same 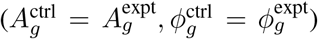, gain 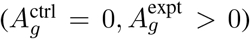, loss 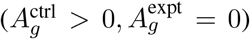, change 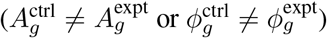, i.e., arrhythmic genes have *A*_*g*_ = 0.

The count for gene *g* at time point *t*_*n*_ under condition *c* was modeled by a negative binomial (NB) random variable:

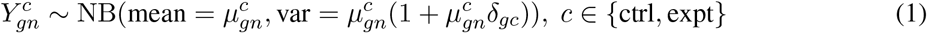

where δ_*gc*_ is the overdispersion parameter that is independent of the time point. For simplicity, we let the dispersion parameter be the same for the two groups, δ_*gc*_ = δ_*g*_.

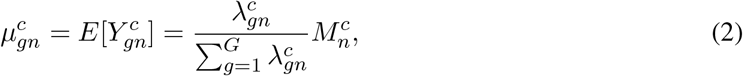

where 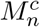 is the sequencing depth for sample *n* in condition *c*, which was defined as 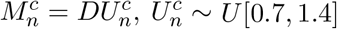 for an nominal study sequencing depth *D*.

For each gene, we drew a mean-dispersion pair (*λ*_*g*_, *δ*_*g*_) from empirical estimates from the multitissue circadian time series data [5]. The relative expression of gene *g* was computed using:

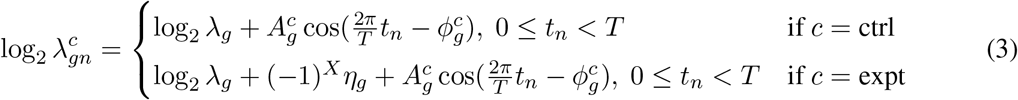

where *X ~* Bern(0.5) determines if a gene is up or down-regulated and rhythm period *T* = 24. We assembled these steps into software packages in R (bharathananth/fauxseq) and Python (bharathananth/pyfauxseq) to encourage future method development and benchmarking of circadian tools.

### Varied Parameters

To limit the number of parameter combinations to be analyzed, we fixed *T* = 24h0, *G* = 10000, *a*_spread_ = 0.5, *η*_min_ = 0.5, *η*_spread_ = 1 and only varied the parameters listed in Table 1 to simulate 4 independent datasets for each combination, resulting in a total of 864 synthetic datasets. A dataset with 6h sampling resolution and 2 replicates per time point will have 24*/*6 *×* 2 = 8 samples.

**Table 1:** Parameters varied in the benchmarking datasets.

| Parameter | Symbol | Qualitative levels | Quantitative levels |
| --- | --- | --- | --- |
| Sampling resolution (time between samples) |  | low, medium, high | 6h, 4h, 2h |
| Replicates per time point |  | low, high | 2, 4 |
| Sequencing depth (nominal per sample) | $D$ | low, medium, high | 5 mill., 20 mill., 50 mill. |
| Fraction of rhythmic genes (in either condition) | $f_R$ | low, medium, high | 0.05, 0.1, 0.2 |
| Minimum amplitude effects | $a_{\text{min}}$ | normal, large | 0.2, 0.7 |
| Fraction of differentially expressed genes | $f_{\text{DE}}$ | absent, present | 0.0, 0.1 |

### Evaluated Methods

We compared 12 different methods from 5 different packages including a simple baseline cosinor regression [1, 6, 7, 8, 9] (Table 2). We used each method under default parameters, both to mimic typical usage and to limit the size of the benchmark, with the false discovery rate (FDR) threshold standardized to 0.05 for the hypothesis testing (HT) methods; the previously established amplitude-based post-hoc filtering (with a threshold of 0.5) within compareRhythms was retained [1]. For methods evaluated using expression-scale input, the same underlying simulated count data were normalized using median-of-ratios normalization and subsequently log_2_-transformed with a pseudocount of 1. Methods evaluated using a count-based input were provided with the raw integer count matrix. Thus, the table describes the input representation used in the present evaluation rather than whether the underlying statistical framework is intrinsically capable of analyzing count data.

**Table 2:** Differential rhythmicity (DR) methods evaluated in this study.

| Package | Method | Framework | Count input | Hypothesis testing (HT) | Model selection (MS) | Version | Source |
| --- | --- | --- | --- | --- | --- | --- | --- |
| *compareRhythms* | CR-cosinor | cosinor regression of normalized data using linear regression (baseline) | ✗ | ✓ | ✗ | 1.5.2 | [1] |
| *compareRhythms* | CR-deseq2 | Negative binomial GLM using DESeq2 [10] | ✓ | ✓ | ✗ | 1.5.2 | [1] |
| *compareRhythms* | CR-dodr | DODR regression [11] | ✗ | ✓ | ✗ | 1.5.2 | [1] |
| *compareRhythms* | CR-edger | Negative binomial GLM using edgeR | ✓ | ✓ | ✗ | 1.5.2 | [1] |
| *compareRhythms* | CR-mod_sel | Linear model selection using BIC. | ✗ | ✗ | ✓ | 1.5.2 | [1] |
| *compareRhythms* | CR-voom | Linear modeling using voom [12] | ✓ | ✓ | ✗ | 1.5.2 | [1] |
| *diffCircaPipeline* | DCP-dPar | Rhythm parameters-based comparison of rhythms | ✗ | ✓ | ✗ | 0.99.0 | [8] |
| *diffCircaPipeline* | DCP-dR2 | $R^2$ fitness-based comparison of rhythms | ✗ | ✓ | ✗ | 0.99.0 | [8] |
| *diffCircadian* | diffCircadian | Likelihood-based testing of rhythm differences | ✗ | ✓ | ✗ | 0.0.0 | [7] |
| *dryR* | dryR | Negative binomial-based model selection | ✓ | ✗ | ✓ | 1.0.0 | [6] |
| *dryR* | dryR-lm | Linear model selection | ✗ | ✗ | ✓ | 1.0.0 | [6] |
| *limorhyde* | limorhyde | Cosinor regression of normalized data using limma [13] | ✗ | ✓ | ✗ | 1.0.1 | [9] |

We formulated the benchmark as a classification task, where each method classified genes into an arrhythmic (AR), same (conserved) rhythms between conditions (R) and differentially rhythmic (DR) classes. All HT methods were structured as two sequential tests: a first hypothesis test to select rhythmic genes and a subsequent test on the rhythmic genes to identify DR genes. The HT methods provided adjusted p-values for each test, which we compared against the chosen FDR threshold. Genes that did not pass the former test were labeled AR and genes that passed both tests were labeled DR. The rest were labeled R. The model selection methods, on the other hand, categorized genes into 5 configurations between the two conditions: arrhythmic, same, gain, loss and change. We pooled all genes in gain, loss and change in the DR class; arrhythmic and same went into AR and R, respectively.

## Performance Metrics

We categorized the ground truth and the predictions from the individual methods into three classes: arrhythmic (AR), rhythmic with the same rhythms between conditions (R) and differentially rhythmic (DR). The latter class includes the “gain”, “loss” and “change” categories in the generated synthetic data (see above). We made this pragmatic choice, since this mapped most closely to the outputs of a majority of methods benchmarked here.

We are therefore dealing with a standard multi-class classification problem—the ground truth for the 3 classes is known and each algorithm classifies genes into the 3 classes. Since the fraction of rhythmic genes observed in most biological samples is small, we also had to address the issue of large imbalance between the AR and (DR, R) classes. At the level of individual classes, *precision* and *recall* are considered more reliable for evaluating performance under class imbalances [14]. These metrics for each class can be aggregated into a single metric termed F_1_ score, which is the harmonic mean between precision and recall. These F_1_ scores can be computed for combinations of classes as simple averages of class F_1_ scores, e.g., for all classes or for only rhythmic gene sets (DR + R). Finally, we quantify the categorization of input data into the 3 classes, accounting for inherent class imbalances, using the multiclass Matthews Correlation Coefficient (MCC) [15].

We dealt with pathological situations in the following manner. For the per class metrics, precision was set to ‘NA’ when a class had no predicted positives, and recall was set to ‘NA’ when a class had no actual positives. F_1_ was set to 0 when either precision or recall was 0, representing a failure state. Otherwise, F_1_ score per class was computed as the harmonic mean of per class precision and recall, with propagation of ‘NA’ values. For the global metrics, means were computed over classes with non-missing values.

## Regressing performance metrics

To evaluate the effect of dataset parameters on performance, we fitted factorial mixed-effects regression models including method, the varied simulation factors, and their interactions with method. Because all methods were evaluated on the same simulated datasets, dataset identity was included as a random effect. To accommodate bounded metrics (Table 3), macro F_1_ was modeled using beta mixed-effects regression with a logit link, while MCC was modeled after Fisher’s *z*-transformation using a linear mixed-effects model. We finally reduced the full model to the best predictive model for its complexity using the Bayesian Information Criterion (BIC).

**Table 3:** Performance metrics of the methods.

| Type | Metric | Range | Interpretation |
| --- | --- | --- | --- |
| Per class | Precision | [0,1] | Fraction of correct calls. Same as (1 - false discovery proportion). |
|  | Recall | [0,1] | Fraction of genes in class discovered. |
| | $F_1$ | [0,1] | Balanced combination of precision and recall. |
| Global | macro $F_1$ | [0,1] | Mean $F_1$ across all classes, i.e., equal weight to all classes. |
| | rhythm $F_1$ | [0,1] | Mean $F_1$ across DR +R classes. |
|  | multiclass MCC | [-1, 1] | Overall performance of classification accounting for class imbalances. |

### Stratification

To gain insight into where different methods succeed or fail, we stratified the benchmark datasets and evaluated performance metrics on these strata. These strata divided the benchmark datasets by:

a. *differential expression (DE) (both true and predicted)*: True DE was available as part of the ground truth. For predicted DE, we detected DE using DESeq2 by fitting a condition-specific cosinor model with a condition-specific MESOR, whose significance we determined at the 0.05 level after multiple testing.
b. *dispersion in gene expression*: Genes were classified into three groups (lower 25% - *low*, 25%-75% - *medium*, upper 25% - *high*) based on empirical dispersion used for data generation. Dispersion is closely associated here with expression level due to the mean-dispersion trend in the empirical data.
c. *differences in circadian parameters* (amplitude and phase) between conditions: Only for genes with true differential rhythmicity, we divided genes independently by amplitude and phase into two groups (lower 50% - *low* and upper 50% - *high*). For amplitudes, we considered absolute differences of log_2_ fold change amplitudes. For phase, we considered the absolute wrapped phase difference, where the phases of arrhythmic genes were assigned randomly. This definition meant that phase differences for loss and gain class genes were less meaningful.

## Benchmarking

We ran the reproducible benchmarking of the methods in the study using Dynamic Statistical Comparisons (DSC) [16] package from the Stephens Lab.

## Results

The goal of differential rhythmicity (DR) analysis is to categorize genes in the dataset into 3 classes: genes that are not rhythmic under any condition (AR class), have conserved rhythms in both conditions (R class), and those that have altered rhythms between conditions (DR class). We benchmarked 12 different DR methods (Table 2) on synthetic RNA-seq benchmark datasets derived from 4 replicates of 216 different parameter configurations with ground truth regarding membership in the 3 rhythm classes (Fig. 1, Table 1).

**Figure 1.**
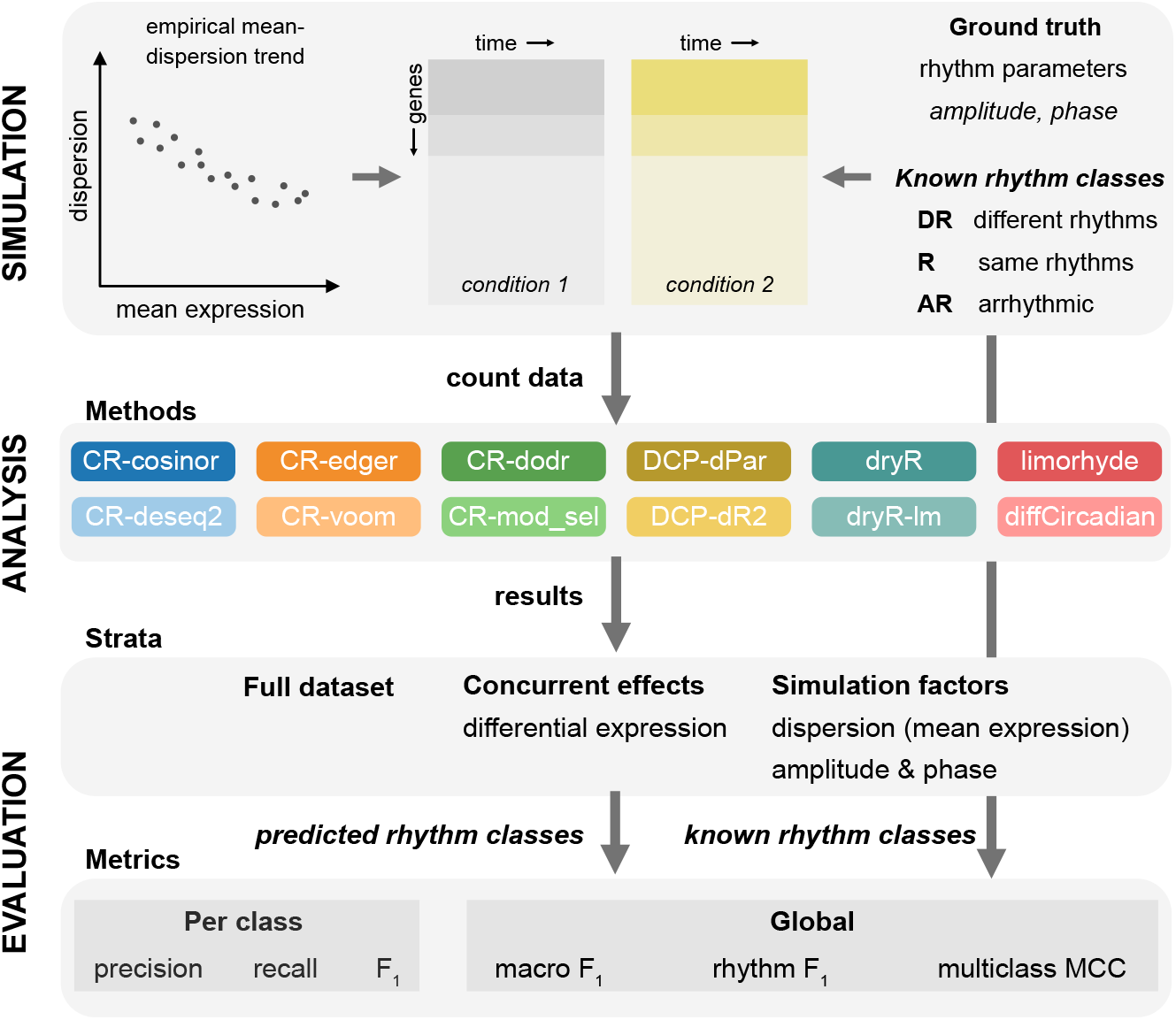
Schematic of the benchmarking.

### A subset of methods consistently performs well overall

We first asked which were the best methods for DR analysis based on overall performance (Fig. 2). We quantified overall performance on the 864 benchmark datasets based on both the correlation between true and predicted class labels (MCC) and the average F1 score across 3 classes (macro F_1_) (Fig. 2A). These two performance metrics were extremely consistent on different dataset-method combinations (Spearman *ρ* = 0.945, *p* < 0.001). The median performance of all methods except CR-mod sel, DCP and diffCircadian was quite similar. However, this boxplot hides the fact that each method has been run on each benchmark dataset resulting in dependence between the underlying data across columns.

**Figure 2.**
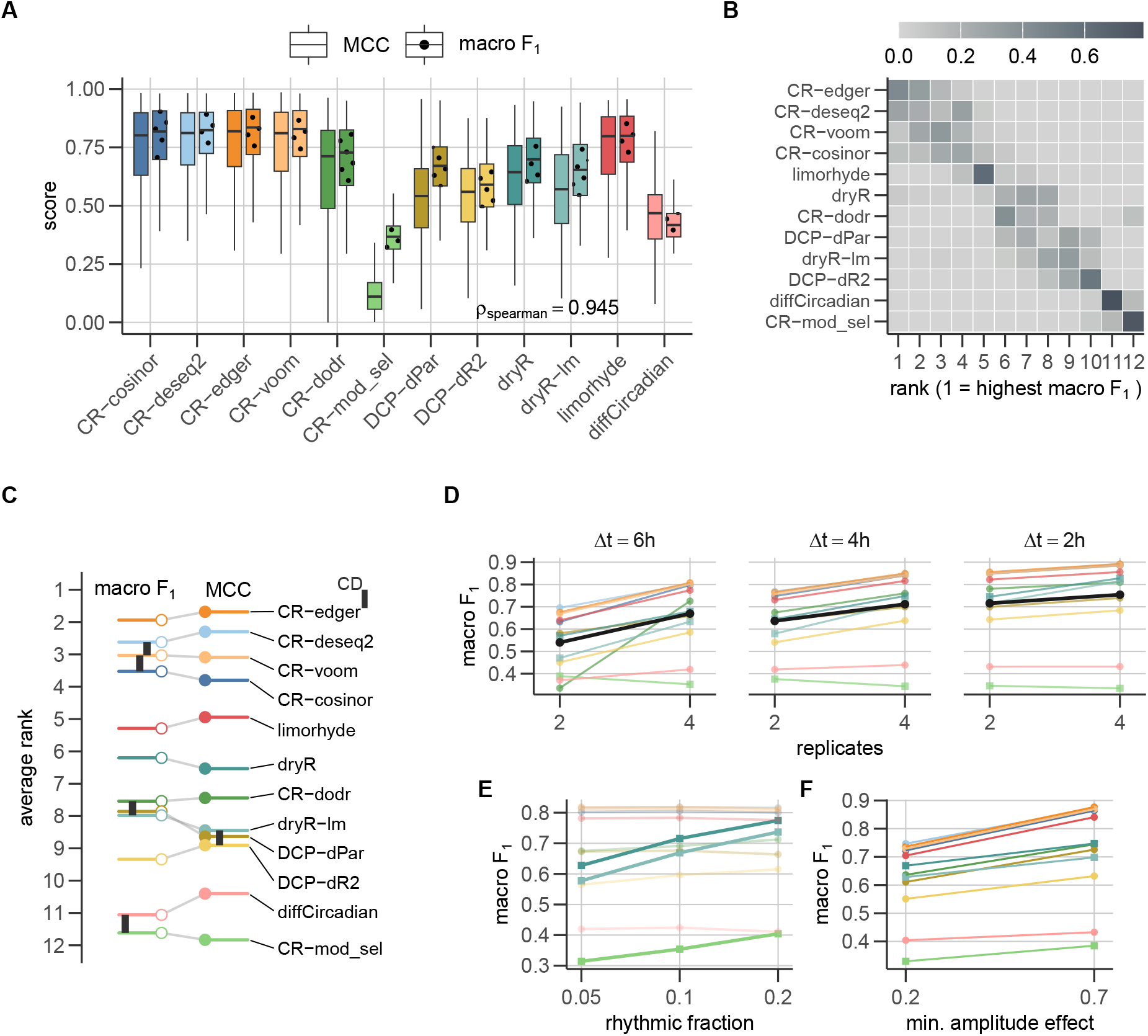
Overall performance of 12 DR methods on synthetic benchmark data. (A) The distribution of multiclass MCC and macro F_1_ scores across the 864 datasets of each method. Spearman correlation between MCC and macro F_1_ across the 864 *×* 12 dataset-method pairs. (B) Distribution of methods’ ranks based on macro F_1_ scores on each benchmark dataset. (C) Comparison of the average ranks of each method (based on both global metrics), including cliques of indistinguishable methods (connected by bars) using a critical difference diagram. (D,E,F) Effects (estimated marginal means) of significant parameters of the benchmark data from beta regression of macro F_1_ scores. Black lines represent averages across methods. Colors for methods are matched across subplots, and hypothesis testing (circle) and model selection (square) methods are shown.

Relative performance of different methods was stable on each dataset. We therefore ranked the 12 methods on each of the 864 datasets using macro F_1_ and computed the distribution of the ranks obtained by each method (Fig. 2B). Five hypothesis testing methods (CR-deseq2, CR-edger, CR-voom, CR-cosinor, limorhyde) and one model selection method (dryR) represented the above median performers, but the CR-dodr surpassed dryR on some datasets. Interestingly, our baseline method (cosinor regression of normalized data) performed better than other bespoke methods. Four of these top 6 methods accept raw count input. Unexpectedly, this conclusion was unaltered when the ranking was performed using MCC as a metric (Fig. S1A). All 12 methods excelled and struggled with the same datasets, as indicated by the strong pairwise Spearman correlation between their MCC and macro F_1_ metrics (Fig. S1B).

We confirmed differences in the method performances across all 864 benchmark datasets using a global Friedman test (*χ*^2^(11) = 7872.62, *p* < 0.001 for macro F_1_ and *χ*^2^(11) = 8032.40, *p* < 0.001 for MCC). We then followed this by a post-hoc Nemenyi test (*α* = 0.05) to establish differences between the average method ranks and indistinguishable method cliques (Fig. 2C). This reinforced our previous conclusion that count-based methods (4 of the top 6) tended to achieve better performance across the benchmark datasets.

We next assessed if and which parameters of the benchmark datasets were determinative of the observed overall metrics. We independently regressed both performance metrics using link-transformed frameworks followed by model reduction using BIC but arrived at very similar models (Fig. 2D-F, Fig. S1D-F). Consistent with expectations, the data reveal that having more replicates, higher sampling resolution, and stronger amplitudes results in better ability to classify genes. Parameters for sequencing depth and for the extent of differential expression (DE) were not retained in the BIC-selected models. We tested whether certain data configurations benefit or are to the detriment of specific methods. The top 6 methods benefited from more replicates with decreasing gain with higher temporal resolution (Fig. 2D). Interestingly, while the hypothesis testing approaches did not vary in performance for different fractions of rhythmic features, model selection approaches profited from a higher fraction of rhythmic features (Fig. 2E). As expected, stronger rhythms (with higher amplitudes) were more easily detected across all methods (Fig. 2F).

### Methods identify differential rhythmicity better than unaltered rhythms

The macro F_1_ is the average of the F_1_ scores for the three different classes (Fig. 3A). All but one method identified arrhythmic genes (AR class) extremely well. The top 6 methods were good at identifying differentially rhythmic (DR class) genes but relatively poor in relation to AR class. Contrary to expectation, many methods performed worst on the rhythmic genes (R class). Across the individual benchmark datasets, methods consistently categorized DR class better than R class genes except for a few poorly performing methods that showed better capability on the R class than the DR class (Fig. S2A).

**Figure 3.**
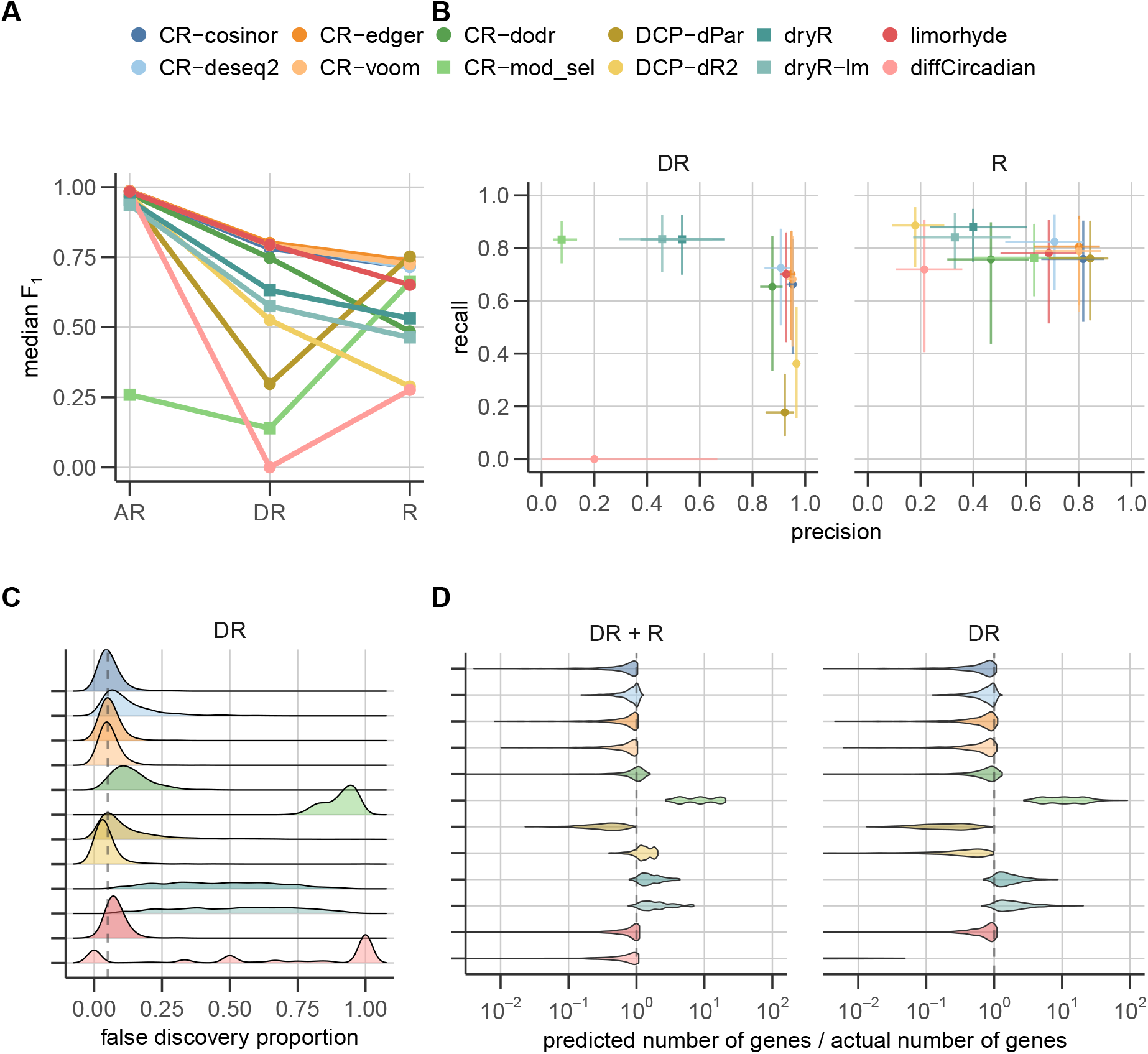
Per class metrics contributing to overall performance. (A) Decomposition of macro F_1_ scores (mean across classes) into F_1_ for each class (summarized as median). Hypothesis testing (circle) and model selection (square) methods are indicated. (B) The median precision and median recall for each method with error bars showing the inter-quantile ranges (25th-75th percentiles) that together contributed to the F_1_ score for the class in (A). (C) The distribution of the observed false discovery proportion for each method. The chosen FDR threshold for the HT methods is shown as a dotted gray line. (D) The ratio of predicted and actual number of genes in total rhythmic (DR + R) and only DR genes (DR).

The per class F_1_ scores are themselves (harmonic) means of precision and recall for the respective class (Fig. 3B). While the hypothesis testing methods (CR-cosinor, CR-deseq2, CR-edger, CR-dodr, limorhyde) achieved high precision in the DR class with relatively poorer recall, the model selection approaches (dryR, dryR-lm, CR-mod sel) provided better recall. On genes with unaltered rhythms (R class), all methods showed middling recall and a broad spread of classification precision. The AR class classification received high precision and recall as expected from the high macro F_1_ scores (Fig. S2B).

We next ascertained how effectively methods controlled false discoveries in relation to the user-defined FDR. (1 - per class precision) is the observed false discovery proportion (FDP), i.e., the fraction of genes assigned to a class that are false discoveries. For HT methods (Table 2), DR calls are generated through two sequential FDR-controlled steps: an initial filtering step to identify genes rhythmic in either condition, followed by a DR hypothesis test among the filtered genes. We therefore compared the empirical FDP of the resulting DR calls with the nominal FDR threshold of 0.05 used in this study. For the DR class, the observed FDP of the HT methods was generally close to the nominal FDR threshold. Among all rhythmic genes (the first of the two sequential steps), the HT methods controlled false discoveries consistently (Fig. S2C). The observed FDP for the AR class across all methods was substantially below the nominal FDR threshold, although these methods are not designed to control AR discoveries. On the other hand, MS methods, which are neither designed to control false discoveries nor guaranteed to do so, suffered from higher fractions of false positives across all three classes (Fig. 3C, Fig. S2C).

### Methods systematically under- or overestimate rhythmic gene set sizes

Although not the intended purpose of DR analysis, studies quantify the number of rhythmic and DR genes and draw biological conclusions about the extent of altered regulation. To evaluate the ability of methods to estimate the size of these gene sets, we computed the ratio of the predicted to true size of gene sets in DR and total rhythmic (DR + R) classes (Fig. 3D). Broadly the trends in both gene sets were similar. The HT methods that belong to the top-performing methods generally and consistently underestimated the gene set sizes, while the MS methods overestimated the gene set sizes. The bottom 4 methods (diffCircadian, DCP-dPar, DCP-dR2, CR-mod sel) identified in Fig. 2C deviated far from the true DR gene set sizes. But the estimate of the total rhythmic set sizes was more accurate even for the bottom 4 methods. Nevertheless, gene set size estimates were unreliable.

### HT methods protect against false discoveries, while MS methods have higher sensitivity to small differences in rhythms

To gain insight into when different methods excelled, we ranked the 12 different methods on different subsets of each benchmark dataset while focusing on the F_1_ score for the DR class (Fig. 4A). We first considered the effect of concurrent differential expression (DE) between the two conditions on the ability of methods to classify different classes. The regression (Fig. 2D-F, Fig. S1D) had previously not retained the DE factor in the BIC-selected models. The ranking of different methods was unaltered on genes with and without DE both based on the ground truth and on the prediction of a DE analysis using DESeq2 [10]. This conclusion was supported by the difference in F_1_ scores for the DR class between the full dataset and each subset (Fig. S3A).

**Figure 4.**
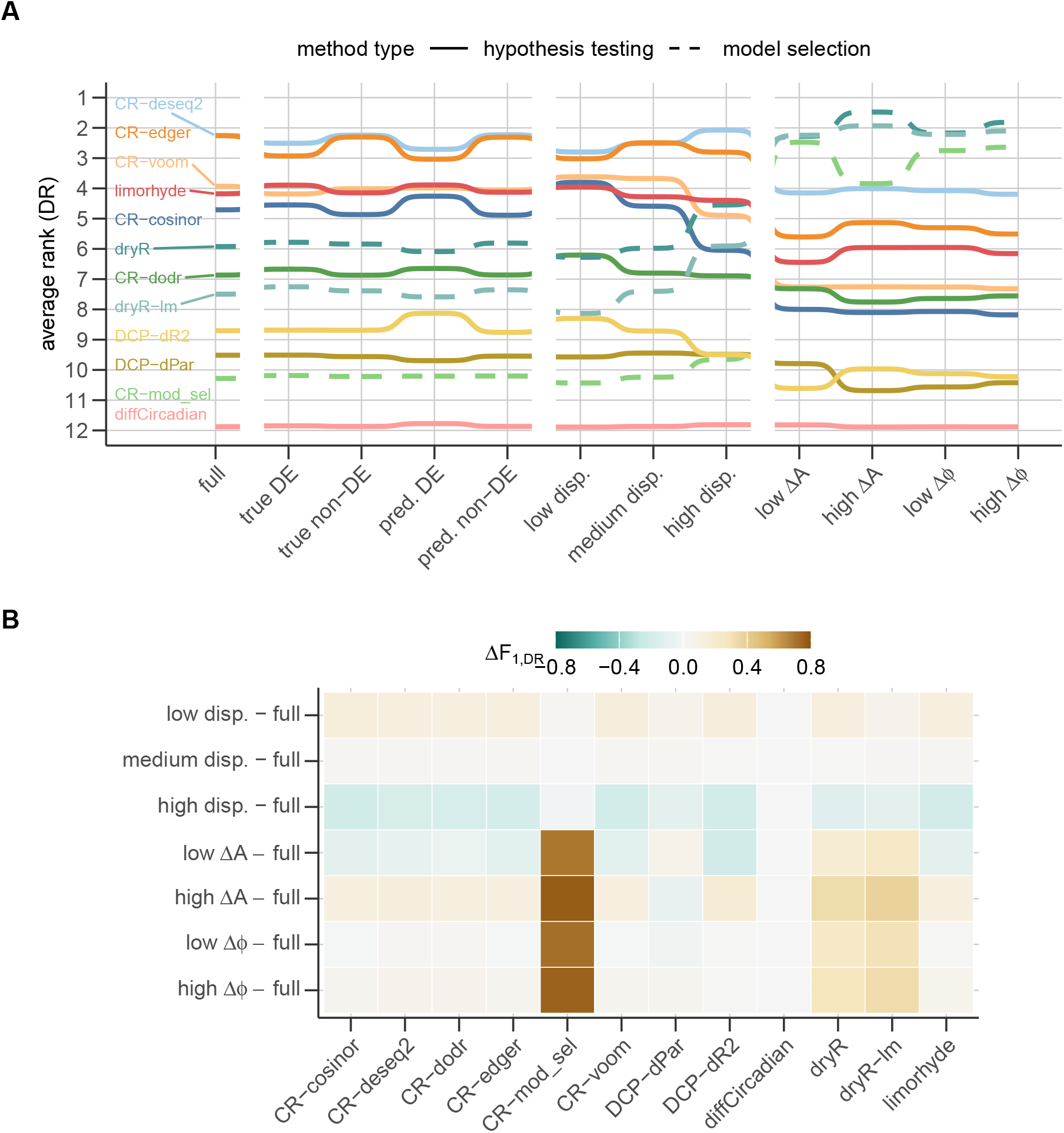
Performance detecting differentially rhythmic genes in different strata. (A) Method ranks based on average F_1_ in the DR class in different subsets of genes within each dataset. (B) Difference in F_1_ score for the DR class of each subset relative to the score for the full dataset.

We next stratified the genes by their dispersion, which is also a surrogate for mean expression due to the mean-dispersion coupling in the data simulation (Fig. 1). Although the ranking remained stable at low and medium dispersion, there was a narrowing of the spread of mean ranks of different methods at the high dispersion (low expression) (Fig. 4A). As expected, all methods performed better among low dispersion and worse among high dispersion genes (Fig. 4B).

We finally tested different methods only on the true DR genes but stratified according to differences in ground-truth rhythm parameters. Surprisingly, the MS methods outperformed the other (HT) methods in this group irrespective of whether the actual differences in circadian parameters (amplitude or phase) were small or large (Fig. 4A). The substantial increase in the average F_1_ for DR class in these subsets for the MS methods helped them achieve top-ranking (Fig. 4B). The F_1_ score in this scenario was driven by the recall (or sensitivity), and it is clear from Fig. 3B that the MS methods achieve high sensitivity. When we expanded our consideration to all rhythmic genes (both same and DR), the qualitative conclusions remained unaltered (Fig. S3B,C).

### Insights are robust to changes in assumptions and thresholds

We next explored the robustness of our observations to thresholds chosen in the analysis (Fig. S4) and assumptions in the synthetic data (Fig. S5). The 9 evaluated hypothesis testing methods used a default FDR threshold of 0.05. We wondered whether any or some methods performed better at a less stringent threshold (of 0.1). However, the same methods scored the best overall based on either metric (Fig. S4A) and correspondingly, the method ranking remained stable (Fig. S4B). The false discovery proportion did not appear to be better calibrated at this threshold (Fig. S4C), and the biases in estimating the total number of rhythmic and DR genes too remained (Fig. S4D).

The empirically estimated mean-dispersion trend made the synthetic data more closely resemble biological RNA-seq data. We decided to assess the effect of this trend by reevaluating metrics after decoupling mean expression and dispersion for each gene (by drawing them independently from empirical estimates). The behavior of individual methods using both global metrics (Fig. S5A) and their relative ranking (Fig. S5B) were stable.

### Rules of thumb for DR analysis

We can now distill our observations into 6 simple rules of thumb for DR analysis of RNA-seq data.

**Rule 1. Prefer methods designed to use raw RNA-seq counts when available**. When raw RNA-seq counts are available, methods that directly accept raw count input should generally be preferred over methods that require normalized and transformed expression as input. In our benchmark, count-based methods represented 4 of 6 most reliable methods across the evaluated scenarios.

**Rule 2. A small group of methods showed consistently strong overall performance, including *compareRhythms, limorhyde*, and *dryR*, depending on the expected fraction of rhythmic genes**. *compareRhythms* (edger, cosinor, deseq2, and voom) and *limorhyde* generally showed high precision and performed particularly well when the fraction of rhythmic genes was low, whereas dryR performed better when a larger fraction of genes was rhythmic. CompareRhythms (mod sel) also showed high sensitivity in scenarios with many rhythmic genes but was prone to more false positives overall. Even the ‘basic’ cosinor regression on normalized data performed well.

**Rule 3. For biomarker discovery or a small set of high-confidence targets, prioritize high-precision methods**. In particular, *compareRhythms* (edger, deseq2, and voom) and *limorhyde* provided a favorable balance for analyses where false positive classifications are of primary concern.

**Rule 4. For functional enrichment analyses, prioritize high-sensitivity methods**. Methods such as *dryR* and *compareRhythms* (mod sel) may be preferable when the primary objective is to recover a larger fraction of truly altered rhythmic genes (even at the cost of some false positives), such as for downstream functional analysis.

**Rule 5. If computationally constrained, opt for *limorhyde* or *compareRhythms* (voom)**. The top 6 best-performing methods, with the exception of *dryR*, were also among the faster methods with limorhyde being the fastest (Fig. S6).

**Rule 6. Do not infer the prevalence of rhythm classes from the number of detected genes**. Most methods showed greater sensitivity for detecting altered rhythms than for identifying genes whose rhythms are unchanged between conditions. Consequently, the numbers of genes assigned to the DR and R classes should not be interpreted independently of method-specific classification performance. Moreover, observed numbers of genes classified as having the same, altered, or absent rhythms should not be interpreted as unbiased estimates of their true proportions and hence should not be used to compare extents of DR across studies.

## Discussion

In this study, we benchmarked 12 different methods for DR analysis arising from 5 different packages on realistic synthetic RNA-seq data with known ground truth. This study was motivated by the numerous questions the author has received at rhythm analysis workshops regarding the ‘correct’ choice of method for DR analysis. DR analysis forms the basis of the controlled-experiment paradigm applied to the study of biological dynamics (in this case rhythms). Effective and appropriate analysis of data arising from such experiments is vital to draw the correct causal conclusions, which have been often flawed [1]. One previous study [3] aimed at providing some clarity on method choice, but their study was limited to biological datasets where ground truth is unknown and must be estimated using other methods. In contrast, we provide simple and easy-to-follow rules of thumb for DR analysis based on the goals of the analysis.

The aim of this benchmarking study was to primarily provide objective directions for DR analysis and not analyze the underlying algorithms. Nevertheless, the surprisingly good performance of the “simple” cosinor regression on the normalized data provided a key insight into hypothesis testing (HT) methods. These methods consist of two sequential steps – a filtering step to find rhythmic genes in either condition followed by DR detection. The top HT methods perform the first step using a single linear hypothesis test, whereas the lower-performing HT methods (CR-dodr, DCP-dR2, DCP-dPar, diffCircadian) perform individual rhythmicity analysis on each condition and then combine the p-values/results, which appears to represent a performance hit. Moreover, the minor role of data properties (mean-dispersion trend, normalized vs. count data) on the ranking implies that the observed differences are more fundamental properties of the algorithms. Finally, all methods struggled in detecting DR in the same datasets suggesting that combining results from different methods is unlikely to yield benefits.

One byproduct of this benchmarking was the development of packages in R and Python (*fauxseq* and *pyfauxseq*) to generate synthetic time series RNA-seq count data with empirically estimated mean-dispersion trends. The methods themselves, however, were largely insensitive to the mean-dispersion trend in the scenarios evaluated, but accounting for the negative-binomial nature of count data appeared to provide count-based methods an advantage over cosinor regression of normalized data. Finally, we implemented the benchmarking using the DSC tool that not only allows easy replication of our study but also effortless addition of newer tools into the benchmark (as they become available). We believe both these features will contribute to the community beyond this specific benchmark.

The results of these studies on synthetic data must be understood within their inherent limitations. Although we included many different data configurations, we had to limit the number of combinations to make the study feasible. We estimated the mean-dispersion trends from 5 different tissues from a single multi-tissue two-condition circadian study [5] and assumed a fixed dispersion across conditions and time points. We also only considered rhythmic expression with a sinusoidal profile, because no DR analysis method, to the best of our knowledge, is currently equipped to detect changes in non-sinusoidal rhythms. As the most common experimental design, we only considered comparison across two conditions. The normalized data used as input to non-count-based methods were derived from median-of-ratios normalization and we did not explore alternate normalization approaches. We did not include circaCompare or RepeatedCircadian, which were included by [3], since the former is not intended for DR analysis according to its author (see github issues) and the latter is designed for longitudinal repeated-measures designs, which we did not explore in this benchmark.

We hope that these simple rules of thumb will make differential rhythmicity analysis more transparent and reproducible, and ultimately help researchers extract more reliable biological insight from time series RNA-seq experiments.

## Acknowledgments

B.A. acknowledges funding from Deutsche Forschungsgemeinschaft (project numbers 511886499 and 528064681). The author also thanks Hanspeter Herzel and Marta del Olmo for useful comments on the draft manuscript.

## Code Availability

The software packages used to generate synthetic RNA-seq data are available in R (10.5281/zenodo.23446285) and Python (10.5281/zenodo.23446234). All the code for reproducing the benchmarking and the bench-marking data underlying the figures is available at 10.5281/zenodo.23475775.

