## Supplemental Figures S1-S6 for "Different, same, or arrhythmic: an empirically informed benchmark of differential rhythmicity methods"

### Supplemental Information

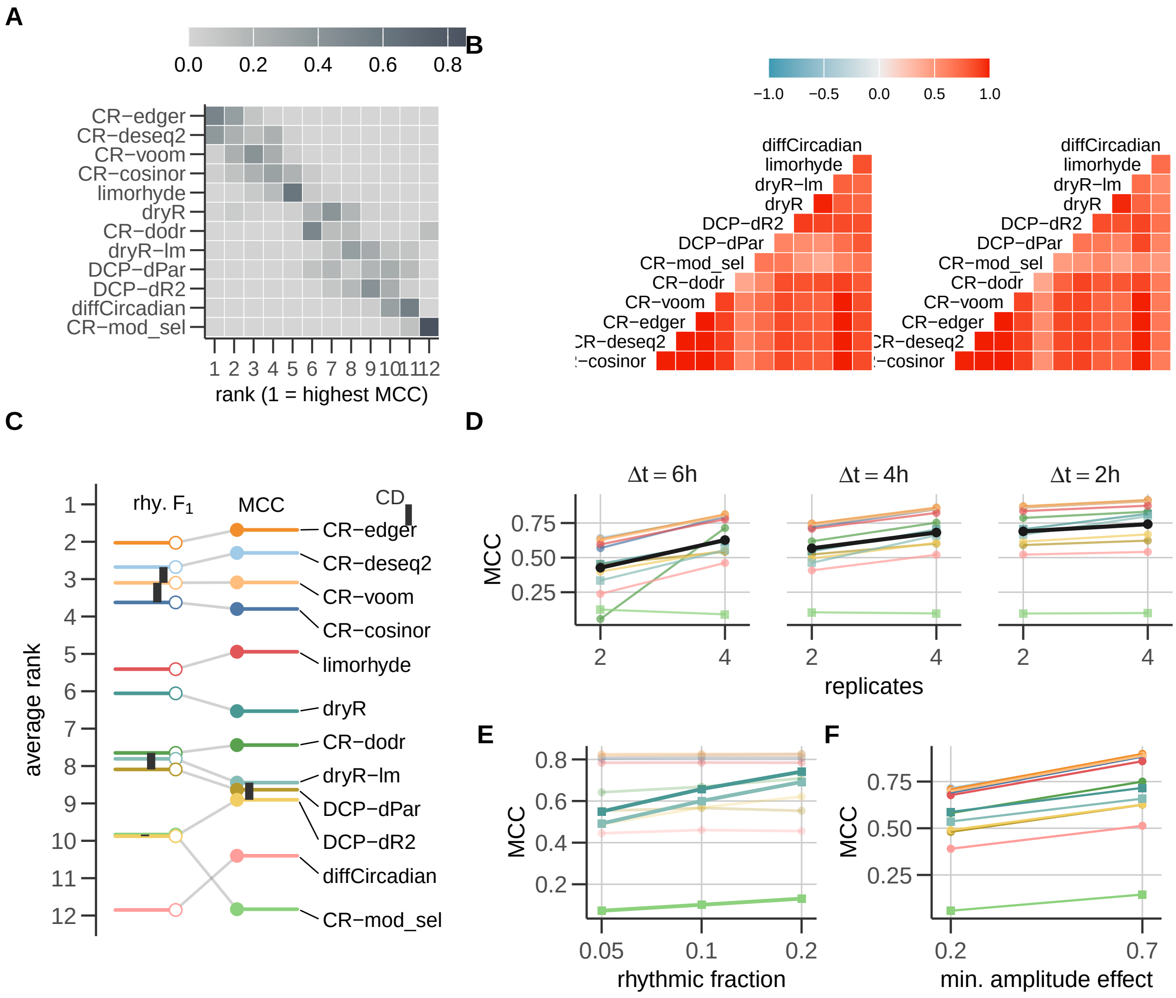


Figure S1: **Overall performance of 12 DR methods on synthetic benchmark data.** (A) Distribution of methods' ranks based on global multiclass MCC on each benchmark dataset. (B) Pairwise correlation between methods on the benchmark datasets based on multiclass MCC (left) and macro $F_1$ scores. (C) Comparison of average ranks of each method (based on both multiclass MCC and average $F_1$ for only rhythmic classes (DR+R)) including cliques of indistinguishable methods (connected by bars) using a critical difference diagram. (D,E,F) Effects (estimated marginal means) of significant parameters of the benchmark data from the atanh-transformed multiclass MCC. Black lines represent averages across methods. Colors for methods are matched across subplots.

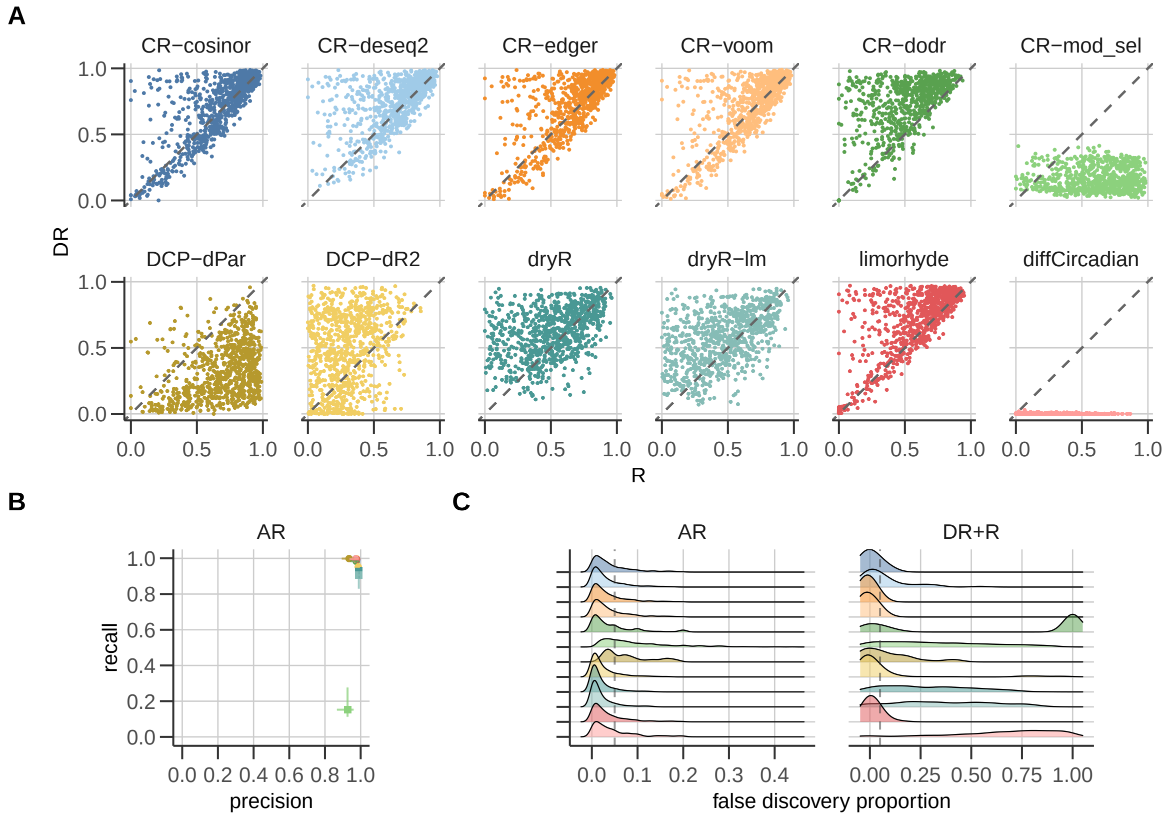


Figure S2: **Per class metrics contributing to overall performance.** (A) Scatter plot of the method $F_1$ scores for the DR and R classes across the entire benchmark dataset (compare with Fig. 3A). The dashed diagonal represents balanced performance in a dataset in both R and DR classes. (B) The median precision and median recall for each method with error bars showing the inter-quantile ranges (25th-75th percentiles) that together contributed to the $F_1$ score for the AR class in Fig. 3A. (C) The distribution of the observed false discovery proportion for AR and total rhythmic (DR + R) classes (related to Fig. 3C). The chosen FDR threshold for the HT methods is shown as a dotted gray line.

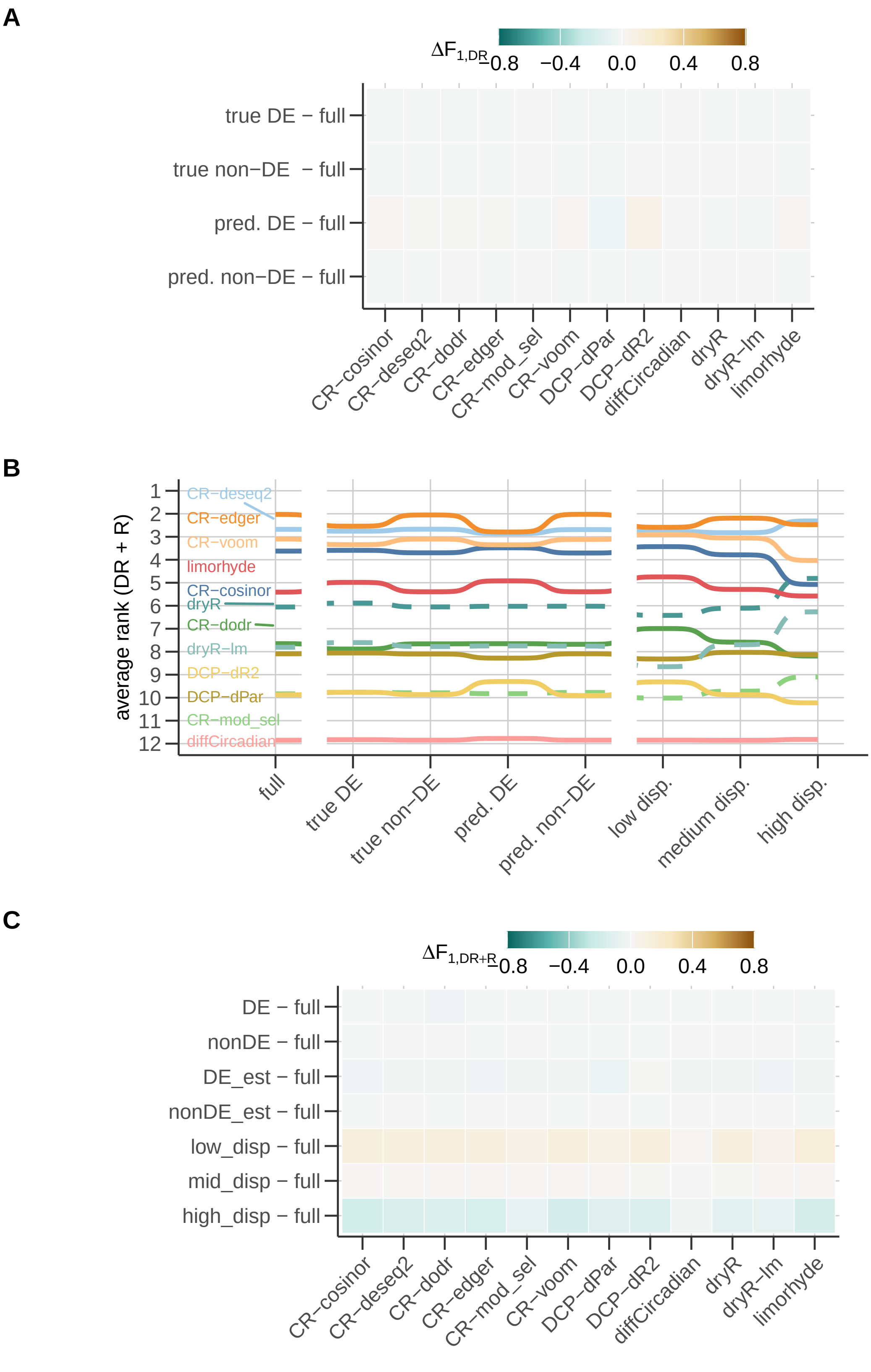


Figure S3: **Performance of detecting rhythmic genes in different strata of genes.** (A) Difference in $F_1$ score for the DR class of subsets related to concurrent differential expression relative to the score for the full dataset. Part of Fig. 4B. (B) Method ranks based on average $F_1$ for all rhythmic genes (DR + R) in different subsets of genes within each dataset. (C) Difference in $F_1$ score for all rhythmic genes (DR + R) for each subset relative to the score for the full dataset.

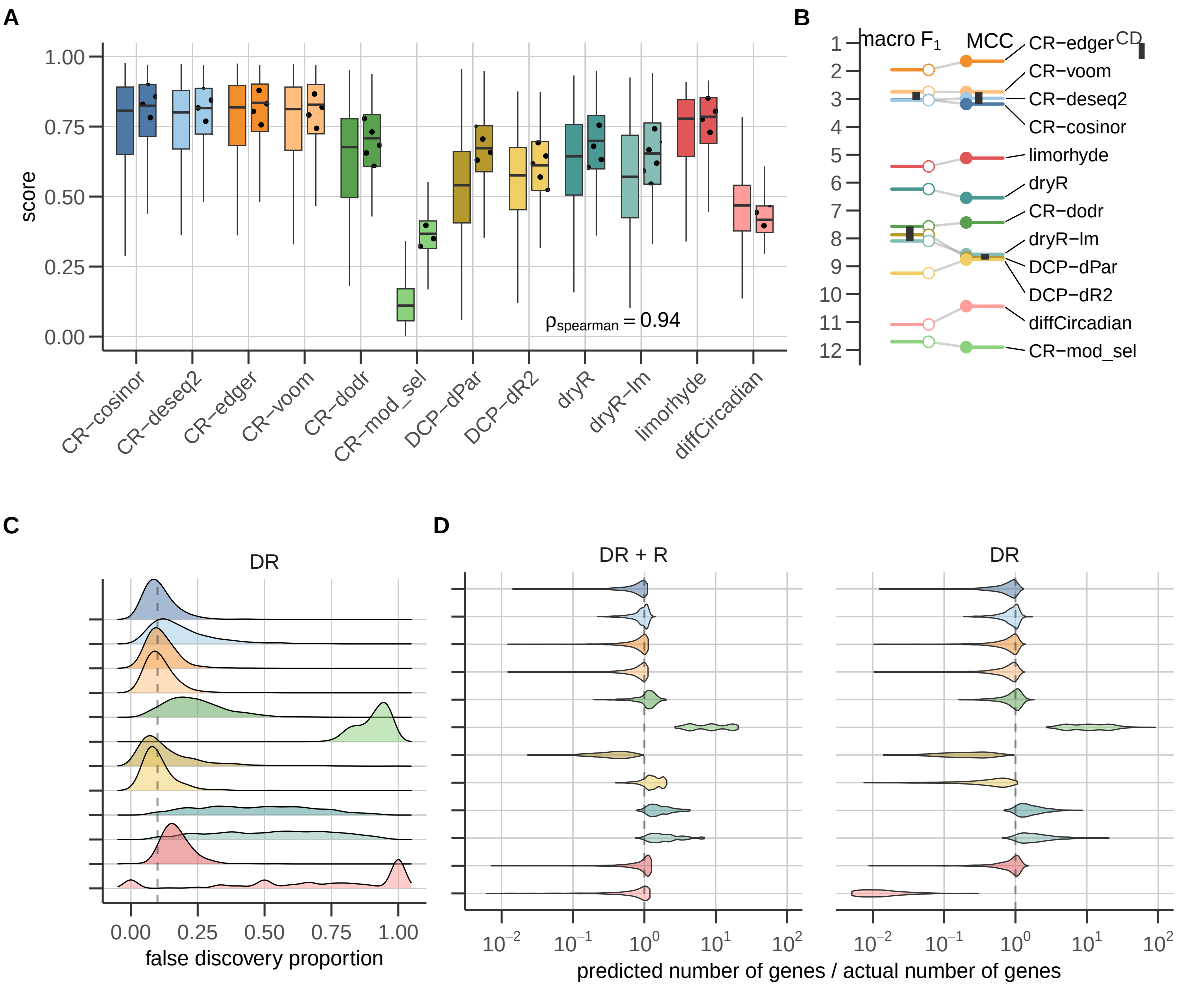


Figure S4: **Overall performance of 12 DR methods at an FDR threshold of 0.1** (A) The distribution of multiclass MCC and macro $F_1$ scores of the methods across the 864 datasets with an FDR threshold of 0.1. Spearman correlation between MCC and macro $F_1$ across the $864 \times 12$ dataset-method pairs. (B) Critical difference diagram showing the average ranks of each method (based on both global metrics) including cliques of indistinguishable methods (connected by bars). (C) The distribution of the observed false discovery proportion for each method. The nominal FDR threshold (here 0.1) for the HT methods is shown as a dotted gray line. (D) The ratio of predicted and actual number of genes in set of all rhythmic genes (DR + R) and only DR rhythmic genes (DR).

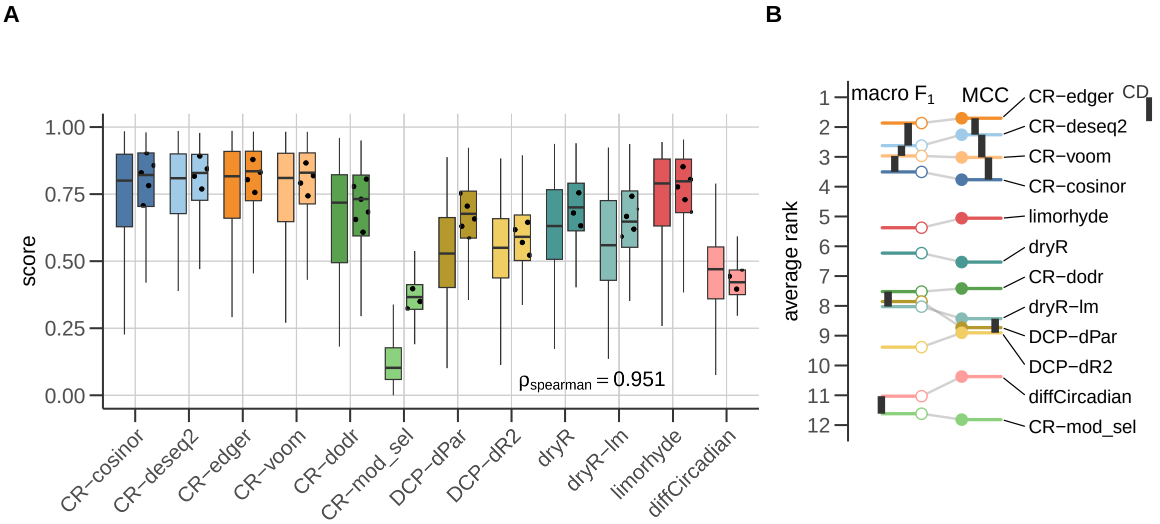


Figure S5: **Overall performance of 12 DR methods on synthetic benchmark data without a mean-dispersion trend.** (A) The distribution of multiclass MCC and macro $F_1$ scores of the methods across the 864 datasets with no mean-dispersion trend. Spearman correlation between MCC and macro $F_1$ across the $864 \times 12$ dataset-method pairs. (B) Comparison of the average ranks of each method (based on both global metrics) including cliques of indistinguishable methods (connected by bars) using a critical difference diagram.

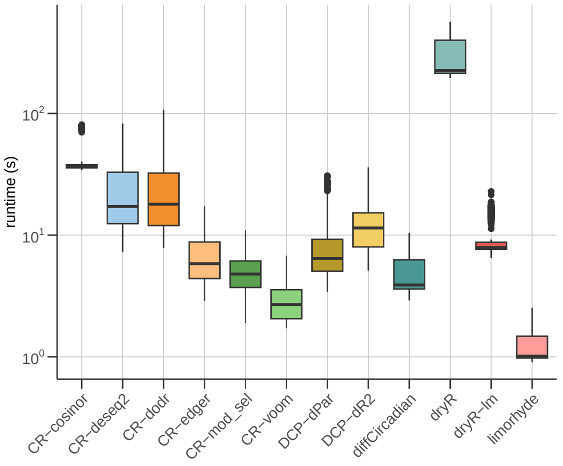


Figure S6: **Run time of different methods on benchmark datasets**
